# Within-host diversity of *Mycobacterium tuberculosis*: weak host association and no transmission signal in a Ghanaian cohort

**DOI:** 10.64898/2026.05.27.727477

**Authors:** Marie Nancy Séraphin, Jane Sandra Afriyie-Mensah, Michael Asare-Baah, Julia Chariker, Charles Domotey, Ernest Kwarteng, Magalie Zoungrana, Sheila Mireku Appah, Honesty Mensah Ganu, Michael Amo Omari

## Abstract

**Background:** Within host diversity (WHD) in *Mycobacterium tuberculosis* is increasingly used to infer transmission and treatment response, but it lies at low allele frequencies where sequencing error and reference mapping artifact can be mistaken for signal.

**Methods:** In a cohort in Accra, Ghana, we measured WHD using deep short read whole genome sequencing and LoFreq with a depth-conditional detection limit and an artifact-aware mask and assessed reproducibility using within-patient biological replicates.

**Findings:** Among 83 single strain infections, about a fifth of called variants were recurrent artifact, and the corrected WHD burden was modest (median 16 variants per patient). WHD was higher in *M. africanum* than in lineage 4 (median 26 versus 14; p = 0.002) and was not explained by host or clinical features. Over the first two months of treatment the within host bacterial population drifted and cleared without detectable positive selection. Patients with an unfavorable two-month outcome carried higher baseline WHD (median 20.5 versus 14.0; p = 0.014). Shared within host variants did not mark transmission, and a single sputum undercaptured diversity.

**Interpretation:** WHD in early treatment is modest, lineage-patterned, and drift-dominated, and reading it for clinical or epidemiological signal depends first on measuring it defensibly.

**Research in context:** *Evidence before this study:* We searched PubMed, Semantic Scholar, and Scopus (via Consensus) through July 2026 for within-host *Mycobacterium tuberculosis* diversity, using terms combining “within-host / minority / intra-host variants”, “sequencing error / artifact”, “transmission”, “treatment outcome”, and “*M. africanum* / West Africa”. Within-host *M. tuberculosis* diversity is real but small and inconsistently measured, and low frequency calls are strongly shaped by sequencing and mapping artifact[1–5]. Shared minority variants have been proposed to sharpen transmission inference[1], while baseline diversity has not predicted treatment outcome[6]. Artifact-aware within-host data that include *M. africanum*, which causes a large share of West African tuberculosis, are lacking[7,8].

*Added value of this study:* Using a depth-conditional empirical limit of detection with a monomorphic site false positive check, we provide an artifact-aware within-host diversity measurement in a Ghanaian cohort: a modest burden (median 16 iSNVs) an *M. africanum* elevation, weak host association, no transmission signal, and marked single-sputum undercapture.

*Implications of all the available evidence:* Within-host diversity holds public health promise as densely captured minority variants could refine transmission chain reconstruction, flag mixed infection, and surface emerging drug resistance earlier than consensus genomes allow. Realizing that promise requires sampling and measurement dense enough to recover the signal that single sputum specimens miss.

## Introduction

Tuberculosis, caused by the *Mycobacterium tuberculosis* complex, remains among the leading causes of death from a single infectious agent worldwide[9]. Whole genome sequencing (WGS) is widely used to compare *M. tuberculosis* strains between patients, typically as a single consensus genome per infection[10]. An infection, however, is not one genotype but a population of bacteria, and deep sequencing resolves the positions in the genome where a minority of that population carries a different allele from the dominant one[11,12]. These low-frequency, or minor, variants (intra-host single nucleotide variants, iSNVs) make up a patient’s within-host diversity (WHD): the standing genetic variation that mutation generates during infection, on which drift and selection then act through disease and treatment[4]. Because the WHD holds this evolving variation, and because it carries signal that a consensus genome discards, WHD is increasingly used to infer recent transmission, to detect emerging drug resistance before it fixes, and as a marker of treatment response[13–15].

The cost of discarding *M. tuberculosis* within-host variation is clearest for transmission inference. Recent transmission is usually inferred from consensus genomes, with clusters of putative recent transmission defined by a genomic distance of 5 to 12 single nucleotide polymorphisms (SNP)[13]. Because WHD in a single patient can be as large as the differences between epidemiologically linked cases[11], a threshold applied to consensus genomes can place linked cases outside a cluster or unlinked cases inside it, leaving contact investigation inefficiently targeted[16,17]. Tracking WHD rather than collapsing it to a consensus has therefore been proposed as a higher resolution way to reconstruct who infected whom [5,17].

Using WHD to reconstruct transmission presupposes that it can be measured reliably. Also, what WHD looks like once it is measured carefully is not well established. Basic questions remain unresolved, such as how large WHD is, once artifact is controlled, whether it tracks host, clinical, or bacterial lineage characteristics at diagnosis, how it changes during treatment and whether it relates to treatment response, and whether shared within-host variants mark person-to-person transmission. The number of specimens needed to capture WHD is likewise unsettled, with direct consequences for how prospective sampling should be designed. All of this turns on the same measurement problem: true minor variation has to be distinguished both from sequencing and sample processing noise and from multi-strain mixed infection, and in the absence of a common, error-aware standard for doing so, reported diversity varies widely across studies[3].

We address this gap in a tuberculosis treatment cohort enrolled at a tertiary hospital in the Greater Accra Region, Ghana, in which each patient contributed biological replicate sputum cultures[18]. We calibrated a depth-conditional detection limit to the per-site sequencing error process[2,19] and use the within-patient replicates to assess reproducibility; against that error-aware detection limit we define WHD from deep sequencing and ask what it looks like at diagnosis, whether it correlates with host, clinical, and bacterial lineage characteristics, how it changes over the first two months of treatment, and whether baseline diversity is associated with the two-month treatment outcome. We then turn to its public health promise, asking whether shared WHD identifies person-to-person transmission and whether collecting a second sputum specimen rather than one meaningfully changes the diversity captured. Our aim is less to add another estimate of WHD than to establish what it is once measured defensibly, and thereby which of its proposed clinical and epidemiological uses its signal can actually support.

## Methods

### Study population and specimens

We studied adults initiating treatment for pulmonary TB at a clinic in Accra, Ghana. At enrollment (baseline, M0) each patient provided two spot sputa and one early morning sputum sample, and a subset with continuous productive cough provided follow-up spot and early morning sputa at one month (M1) and two months (M2) of treatment. Clinical and demographic data, including the two-month treatment outcome, were recorded for all participants.

### Sequencing and read processing

Genomic DNA was extracted from sputum samples decontaminated with 1% final concentration of sodium hydroxide and cultured for up to six weeks in mycobacterial growth indicator tubes (MGIT). Samples were sequenced using Illumina short read NovaSeq platform to high depth (median per-specimen depth in the analytic set 284-fold). Sequencing reads were processed using a previously reported pipeline[1,2] implemented in Nextflow[20]. Briefly, reads were mapped with BWA-MEM[21] and processed with SAMtools[22]. To avoid inflating minor variant calls in the more divergent lineages[23], reads were mapped to the reconstructed most recent common ancestor genome of the *M. tuberculosis* complex on H37Rv coordinates[24,25]. Lineage and mixed infection status were assigned with TB-Profiler[26,27]; a specimen carrying more than one major lineage call (a “;” in the TB-Profiler barcode) was flagged as mixed. Genomes were required to pass a minimum depth quality control of 50-fold.

### Within-host variant calling and the depth-conditional detection limit

Low-frequency variants were called with LoFreq[28], a sequence-quality-aware caller that models per-run, per-position error and is conservative in *M. tuberculosis* WGS, with a single nucleotide detection limit near 3% minor allele frequency (MAF)[29]. Insertions and deletions and the PE/PPE/PE-PGRS gene family, which are repetitive and prone to short read mapping artifact were removed[1,2], and positions of extreme depth were trimmed[1]. Analysis was restricted to biallelic single-nucleotide positions in the candidate MAF band between 0.02 and 0.50 in genomes passing the 50-fold depth quality control, and each candidate position was additionally required to have at least 50-fold local depth. Rather than impose a single flat MAF or read-count threshold, we set a depth-conditional limit of detection empirically from the sequencing error observed at the cohort’s own monomorphic positions (below the 2% band) at which a minor allele reflects error rather than true within-host variation[2,30]. Within depth strata (50-100, 100-150, 150-200, 200-300, 300-600, and >600-fold), the detection floor a(DP) was the smallest alternate read count exceeded by no more than a target fraction *q = 0.01* of monomorphic positions; the floor is therefore the empirical upper tail of the error process at each depth. The adopted floors were 2, 3, 3, 4, 5, and 34 alternate reads across the depth strata, corresponding to detectable minor allele frequencies of 0.026 down to 0.014 and rising again to 0.020 in the deepest stratum, and the realized false-positive rate at each floor was at or below the 0.01 target in every stratum (**Table S1b, Figure S1**). Because the floor is read directly from the empirical error tail, it rises with depth as that tail fattens and requires no distributional assumption. A candidate variant was retained if its alternate read support met a(DP) for its depth and we set a conservative strand bias filter requiring the weaker strand to carry at least 10% of the minor allele reads[2]. The full low-frequency variant detection procedure and its calibrated constants are given in **Table S1a**, and the depth conditional limit in **Table S1b**. All frequencies are reported as folded MAF (equivalently, alternate allele frequency in [0.02, 0.98]); trajectories over the two month follow-up period were additionally computed in signed alternate allele frequency (below).

### Reproducibility of calls

Reproducibility was quantified on the final analytic calls as Lin’s concordance correlation coefficient (ccc)[31] between the paired MAF of within-patient biological replicates: spot versus spot (n = 46 pairs) and spot versus early morning (n = 95 pairs). Bootstrap confidence intervals were obtained by resampling patients.

### Candidate band recurrence mask

Standard artifact masks target positions recurrent below the 2% floor and did not remove a class of position that recurred across unrelated patients at 10% to 40% MAF and were dispersed across several hundred genes, consistent with systematic reference mapping discordance rather than shared biology[1]. We therefore masked positions that were recurrent in at least 10 patients in the candidate band, excluding resistance associated genes from masking so that any drug-driven signal would be retained[32]. The 45 masked positions are listed in **Table S3**. Sensitivity to the recurrence threshold was assessed at 3, 5, and 10 patients.

### Cluster-dominated specimen exclusion

Genuine within-host variants are expected to be distributed approximately independently across the genome, whereas localized mapping artifact produces dense clusters of calls within a few base pairs (bp) of one another[1,2]. For each specimen we computed the fraction of calls within 20 bp of a neighboring call (frac_clust) and the pre-de-clustering call count (n_pre) and removed the clustered positions from all specimens (proximity de-clustering). A specimen was additionally flagged as cluster-dominated, and excluded, if frac_clust was at least 0.80 and n_pre was at least 50, that is, if its calls were overwhelmingly local-cluster artifact [1,2].

### Single-strain analytic cohort

The analytic cohort was restricted to single-strain infections. Patients with any cross-lineage mixed-infection specimen were excluded at the patient level; among the 97 sequencing-QC-passed patients, six were excluded on this basis (nine patients were cross-lineage mixed overall, three of whom had already failed sequencing QC). Per-patient within-host burden was then computed from one representative baseline specimen per patient, selected among non-cluster-dominated M0 spot or early-morning specimens; nine patients whose every baseline specimen was cluster-dominated were thereby excluded. The resulting single-strain analytic cohort comprised 82 patients (184 baseline specimens; **Figure 1a**).

**Figure 1.**
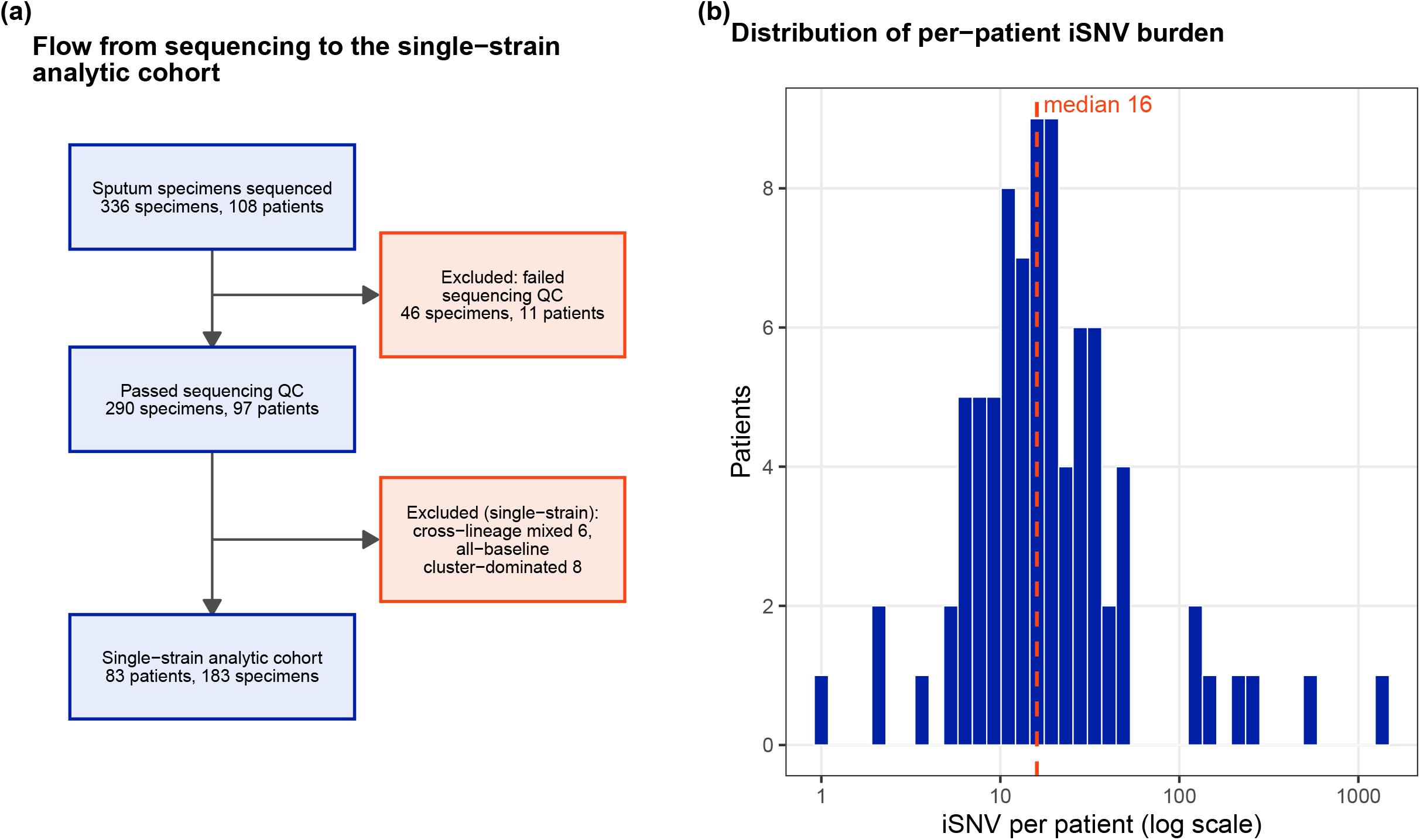
Sample flow and per-patient within-host iSNV burden. (a) Flow from sequencing to the single-strain analytic cohort. (b) Per-patient iSNV counts (log scale); dashed line marks the median (16); detectable in 100.0% of patients.

### Within-host burden and lineage

Per-patient within-host burden was the number of retained minor variants in the representative baseline specimen. Major lineage was grouped as L4, *M. africanum* (L5 or L6), and other-non-L4 (L1). The lineage contrast was tested by Wilcoxon rank-sum (L4 versus *africanum*) and Kruskal-Wallis across the three groups. To evaluate whether the *africanum* elevation reflected reference-mapping artifact, *africanum*-restricted and L4-restricted calls were compared on strand balance, near-floor fraction, repeat-region proximity, and gene concentration, and a sample size-matched bootstrap compared positional recurrence between lineages.

### Host feature model

The association of baseline host and clinical features with WHD was estimated with a negative-binomial regression (MASS::glm.nb)[33]) of minor variant count on HIV status, age, sex, body mass index (BMI), hemoptysis, and per-specimen sequencing depth (per 100-fold). Given the small sample and a severe count outlier, profile-likelihood and nonparametric patient-bootstrap confidence intervals were reported alongside Wald intervals, and the model was refit after excluding far-outlier WHD burden (above the third quartile plus three times the interquartile range). Discrimination was summarized by McFadden pseudo-R-squared and a cross-validated Spearman correlation between predicted and observed burden.

### Longitudinal analysis

For patients with follow-up sputa, one representative specimen per visit was used, a spot specimen where available and otherwise the early morning specimen, and WHD was compared across visits by paired signed-rank tests. A minor variant site was identified by its genomic coordinate together with its reference and alternate base, among the calls retained after depth-conditional detection limit and the candidate band mask. Between two visits the per-patient site sets were partitioned by membership: a site was persistent if it was called at both visits, lost if it was called at the earlier visit but not the later, and emergent if it was called at the later visit but not the earlier. As a sensitivity, the earlier visit set was restricted to sites at a minor allele frequency of at least 0.05. Frequency trajectories of persistent sites were computed in signed alternate-allele frequency, because folded MAF collapses sweeps that cross 0.5. Because a site can leave the called minor variant set either by falling below the detection floor or by rising above the minor allele ceiling to fixation, lost sites were classified by their alternate-allele frequency at the later visit as cleared (below 0.02), intermediate, or fixed (at or above 0.95), and fixed sites were annotated by gene and resistance status.

### Two-month treatment outcome analysis

The two-month treatment outcome was dichotomized as favorable (documented transition to the continuation phase[32]) versus unfavorable (treatment failure, death, still-intensive-phase, second-line or stopped treatment, or loss to follow-up). The primary analysis was intention-to-treat over all patients, counting loss to follow-up (missed month two visit) as unfavorable; a complete-case analysis excluding patients without a recorded outcome was a sensitivity, as was a stricter definition additionally requiring a negative month-two smear. The exposure was baseline WHD. The association was tested by Wilcoxon rank-sum of WHD by outcome and by logistic regression of favorable outcome on the standardized log WHD (odds ratio per standard deviation), with a Firth penalized fit[34,35] for the small event count and a far-outlier-fenced refit.

### Sampling and transmission analyses

Sputum sampling analyses used baseline (M0) spot and early morning specimens from single strain patients passing the 50-fold depth QC (n = 97). Per specimen yield, the number of retained minor variants in the specimen, was compared between specimen types across all specimens by Wilcoxon rank-sum and within patients contributing both types by Wilcoxon signed-rank on per-patient means. Complementarity of specimens was assessed among patients with both a spot and an early morning specimen. For each spot and early morning pair we took the combined set of retained sites and recorded the fraction contributed by the spot alone, by the early morning specimen alone, and by both, averaged these fractions per patient, and compared spot and early morning recovery by paired Wilcoxon.

For transmission, patients were separated by consensus SNP distance, the number of fixed differences between their genomes (biallelic, quality-passing, non-PE/PPE sites), and grouped into clusters at distances of 5, 10, and 12 SNP (12-SNP the primary threshold[36]). A within-host variant was called rare if it was carried by no more than *k* patients, from a cohort prevalence computed once across all 83 baseline patients, with *k* set at 2, 3, 5, and 8 (the last corresponding to 10% of the baseline patients). Two patients of the same major lineage were linked if they shared at least one rare within host variants[1], computed first from one specimen per patient and then from two (spot and early morning). We then tested whether these links were transmission-specific: the fraction of linked pairs also within 12 SNP and within 2 km of residence (from GPS coordinates[18]) was compared against chance and residential distance was compared between linked and unlinked pairs of the same lineage by Mann-Whitney.

## Results

We analyzed data from 108 adults with culture-confirmed pulmonary tuberculosis enrolled at diagnosis in Accra, Ghana, collected multiple baseline sputum specimens per patient (two spot and one early morning), and sequenced *M. tuberculosis* to high depth from 336 specimens. After sequencing quality control (290 specimens from 97 patients) and restriction to single strain infections, the analytic cohort comprised 83 patients, each represented by one baseline specimen for the burden analyses (**Figure 1a**). In this cohort the median age was 39 years (IQR 30 – 52), 77.1% were male, with a median BMI of 18.0 (IQR 15.9 – 19.9), and 7.2% were living with HIV (**Table 1**). Lineage 4 accounted for 68 patients (81.9%) and *M. africanum* (L5 or L6) for 13 (15.7%).

**Table 1.** Characteristics of the single strain analytic cohort.

| **Characteristic** | **Summary** |
| --- | --- |
| Patients (N) | 83 |
| Age (years; median (IQR)) | 39.0 (30.0 to 52.0) |
| Age < 35 years | 29 (34.9%) |
| Age 35 – 54 years | 35 (42.2%) |
| Age 55 years + | 19 (22.9%) |
| Female Sex | 19 (22.9%) |
| Male Sex | 64 (77.1%) |
| HIV- | 77 (92.8%) |
| HIV+ | 6 (7.2%) |
| BMI median (IQR) | 18.0 (15.9 to 19.9) |
| BMI <18.5 | 45 (54.2%) |
| BMI 18.5-24.9 | 34 (41.0%) |
| BMI 25+ | 4 (4.8%) |
| Lineage *M. africanum* | 13 (15.7%) |
| Lineage L4 | 68 (81.9%) |
| Lineage non-L4 | 2 (2.4%) |
| Lineage L1 | 2 (2.4%) |
| Lineage L4 | 68 (81.9%) |
| Lineage L5 | 4 (4.8%) |
| Lineage L6 | 9 (10.8%) |
| Hemoptysis at diagnosis | 23 (27.7%) |
| Weight loss at diagnosis | 80 (96.4%) |
| Fever at diagnosis | 54 (65.1%) |
| Night sweats at diagnosis | 35 (42.2%) |

Against the depth-conditional detection limit (**Figure S2a**), the final within-host calls were moderately reproducible across within-patient biological replicates, with Lin’s concordance correlation of 0.55 for spot versus spot (n = 46 pairs) and 0.57 spot versus early morning pairs (n = 95 pairs; **Figure 2b**). That the two pairings were indistinguishable is informative: once recurrent artifacts were removed, a single spot and a single early morning specimen are equivalent, and the residual between-specimen variation reflects stochastic sampling of the same bacterial population rather than compartment-specific difference. A candidate band mask, which removes above floor positions recurrent across unrelated patients that conventional sub-2% filters retain, removed roughly one third of candidate calls and approximately halved the median per-patient burden (**Figure S2b**). In the final cohort the median WHD was modest, 16 minor variants per patient (IQR 10-27, range 1 to 1291), detectable in every patient (**Figure 1b**). Two specimen restrictions completed the single-strain analytic cohort. Nine patients contributing 16 samples carried cross-lineage mixed infections, which produced an extreme per-patient burdens of 1700 to 2300 minor variants (**Figure 2a**); because lineage mixing inflates apparent within-host diversity independent of within-host evolution[6], these patients were excluded, six of them among the sequencing QC passed set shown in **Figure 1a** (the 16 mixed-infection specimens are listed in **Table S2**). Separately, nine patients whose every baseline specimen was cluster-dominated, the signature of localized mapping artifact rather than dispersed within host variation, were excluded. Several excluded patients were non-L4, so both restrictions are conservative with respect to the lineage contrast below.

**Figure 2.**
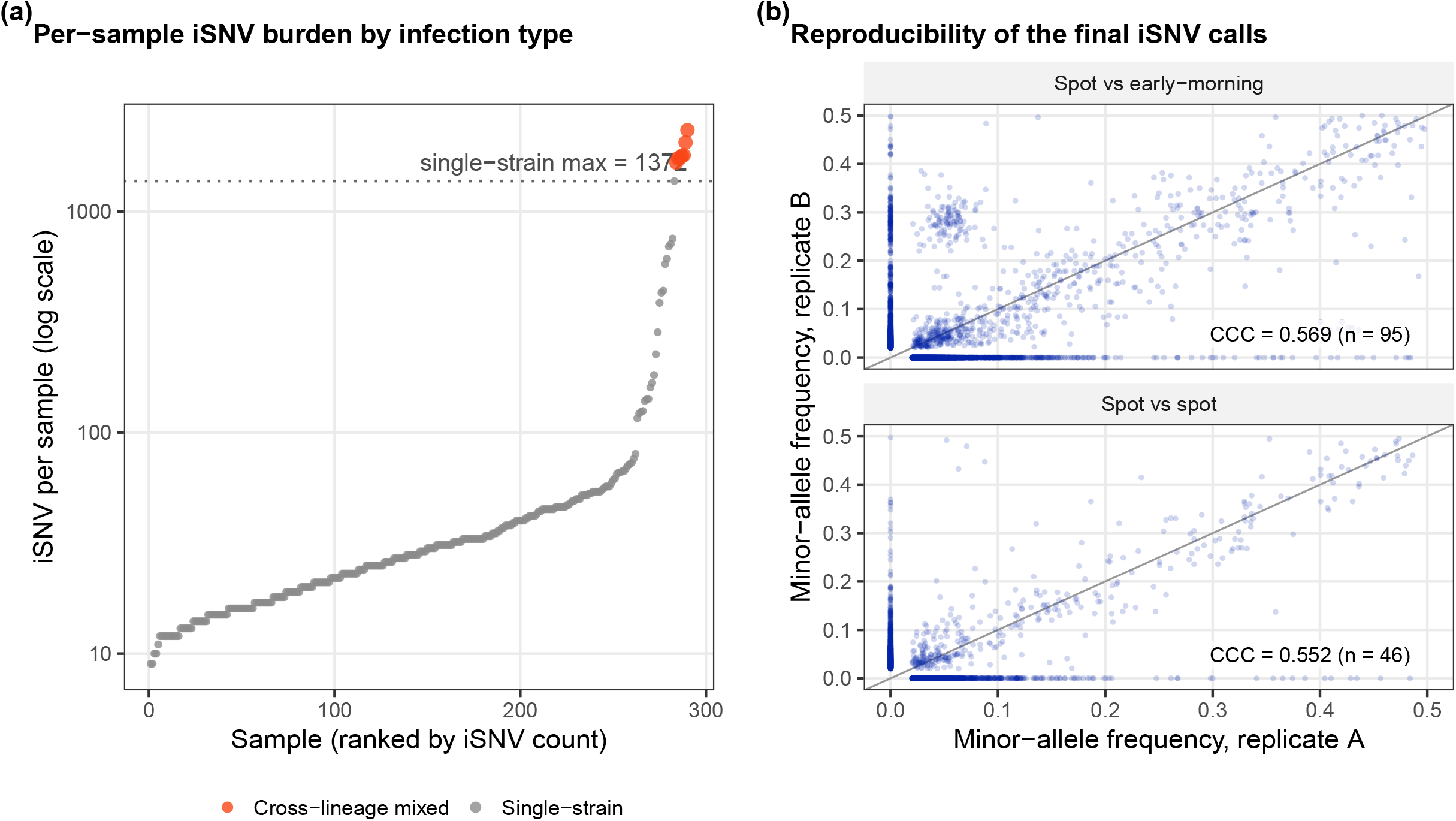
Within-host iSNV burden by infection type, with replicate reproducibility. (a) Per-ample iSNV counts ranked (log scale); cross-lineage mixed infections (orange) form the extreme-burden tail, and the per-sample maximum falls from 2332 to 1372 once they are set aside (single-strain). (b) Paired minor-allele frequencies for spot-spot (n = 46) and spot-early-morning (n = 95) biological replicates, Lin’s concordance correlation annotated. Reproducibility is moderate and, indistinguishable between spot-spot and spot-early-morning: once recurrent artifacts are removed, a single spot and a single early-morning specimen are equivalent, with no compartment-driven sampling difference.

### Within-host diversity is elevated in *M. africanum*

Within-host burden was higher in *M. africanum* than in L4: median 26 iSNV per patient (IQR 21 – 41, n =13) versus 14 (IQR 9-24, n = 67) (Wilcoxon p = 0.0016; three-way Kruskal-Wallis across L4, *africanum*, and L1 p = 0.0036; **Figure 3a**). Because *M. africanum* is the lineage most divergent from an L4-derived reference[24], we tested whether reference bias could account for the difference. The contrast persisted at every masking threshold, and *africanum*-restricted calls were cleaner than L4-restricted calls on strand balance, near-floor fraction, and repeat-region clustering, the opposite of the expectation under lineage-specific reference bias. A size-matched recurrence bootstrap placed the *africanum* value at the upper edge of the L4 distribution but not significantly beyond it (empirical p = 0.12; **Figure S3**). We interpret the elevation as predominantly biological, with a residual confined to paralog and repeat loci (*mmpL5, Rv2081c, pknH*).

**Figure 3.**
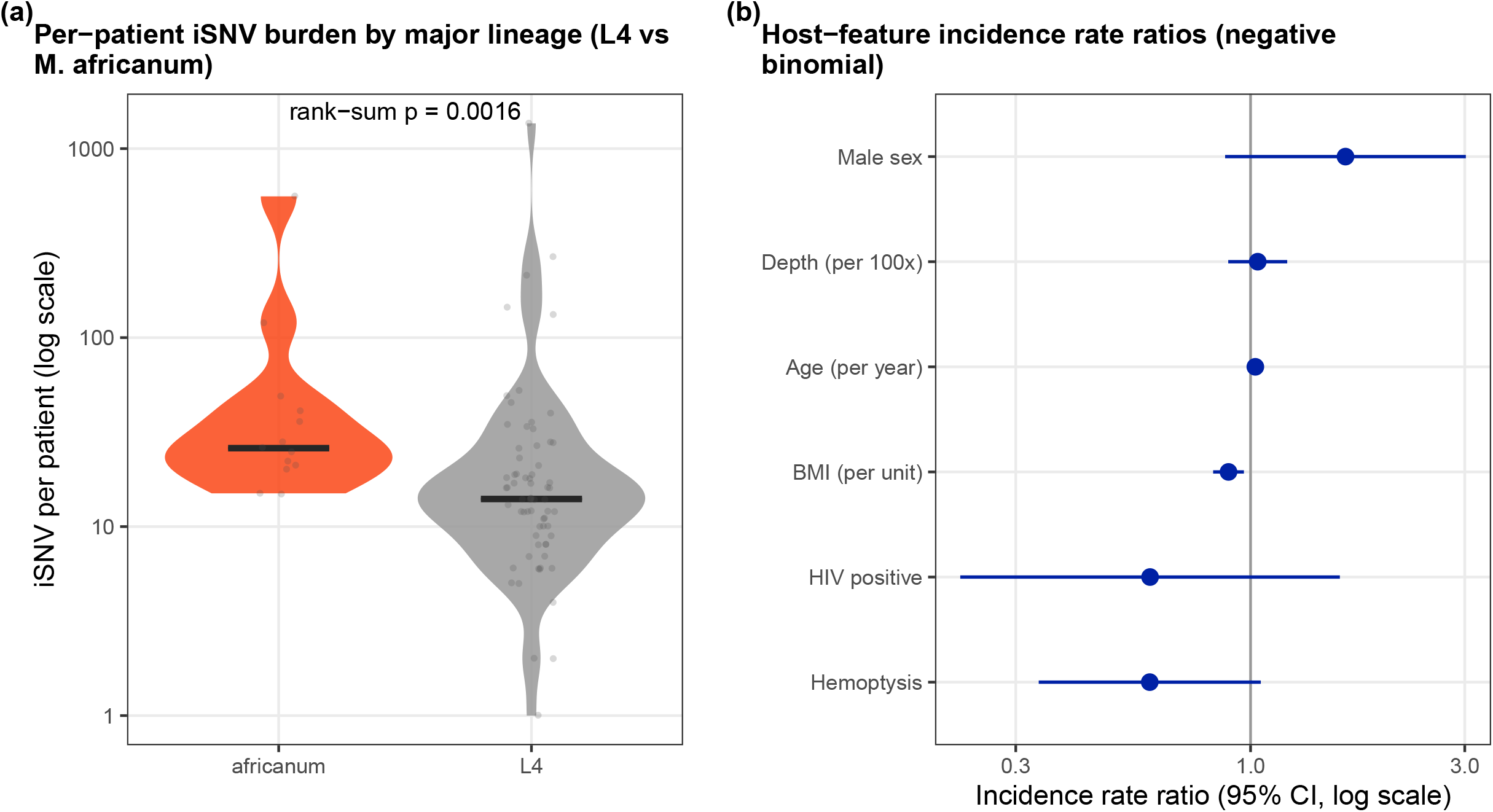
Lineage patterning and host prediction. (a) Per-patient iSNV burden L4 versus *M. africanum* (L5/L6; two L1 patients excluded); africanum higher (rank-sum p = 0.0016). (b) Negative-binomial incidence rate ratios for baseline host features (95% CI, log scale); cross-validated ranking weak (Spearman rho = 0.225).

Host factors tested did not robustly explain WHD. HIV status, age, and BMI were nominally associated with WHD in a negative-binomial model but did not persist after fencing far-outlier burdens (fenced age p = 0.30, BMI p = 0.11, HIV p = 0.68). Read depth was not associated with burden, and a cross-validated ranking was weak [Spearman *rho* 0.23 (95% CI: 0.14 – 0.27); **Figure 3b**). We report the fenced result as primary and the nominal associations as outlier driven.

### Within-host diversity drifts and clears during early treatment

In patients with follow-up sputa, WHD was stable across the first two months of treatment (medians 16, 18, 17 at M0, M1, and M2; paired tests not significant), while the underlying variant composition turned over: between M0 and M1 a median of 2 sites persisted per patient, 10 were lost, and 12 emerged (**Figure S4a, S4b**). Lost sites cleared to an alternate frequency below 0.02 in 96% of cases (M0 to M1) and 92% (M0 to M2), with a small tail that sweep to fixation (2.3% of lost sites by M1 and 6.7% by M2; **Figure S4c**). No fixation occurred in a resistance-associated gene, and the fixations were clonal, concentrated in two patients, indicating compartment or subclone replacement rather than independent selection. We interpret the early within-host dynamics as drift and clearance without detectable positive selection.

### Higher baseline diversity is associated with unfavorable 2-month outcome

We tested whether baseline WHD was associated with the two-month treatment outcome, coded as favorable (documented transition to the continuation phase) or unfavorable; the motivation is the growing use of WHD as a marker of treatment response[14,15]. Ten patients lacked a month-2 outcome; all had missed the month-2 visit and exited follow-up early, and their baseline diversity was higher than the favorable group, indicating informative missingness. We therefore analyzed the outcome by intention-to-treat across all 82 patients, counting loss to follow-up as unfavorable, which gave 59 favorable and 24 unfavorable outcomes, and retained a complete case analysis as a sensitivity.

Baseline WHD was higher in patients with an unfavorable outcome (median 20.5 versus 14.0 iSNV; Wilcoxon p = 0.014; **Figure 4a**). The association measured by logistic regression was directionally consistent but not significant in the full sample (odds ratio 0.66 per standard deviation of log diversity for a favorable outcome, 95% CI 0.40 – 1.06; Firth-penalized 0.68) and strengthened after fencing far-outlier burdens (odds ratio 0.43, p = 0.0092; **Figure 4b**), indicating that the association was not driven by extreme burden patients. The complete case analysis attenuated to a trend, consistent with the excluded losses to follow-up carrying signal, and a stricter definition additionally requiring a negative month-2 smear was null (odds ratio 1.00), indicating that the association tracks clinical progression rather than smear conversion.

**Figure 4.**
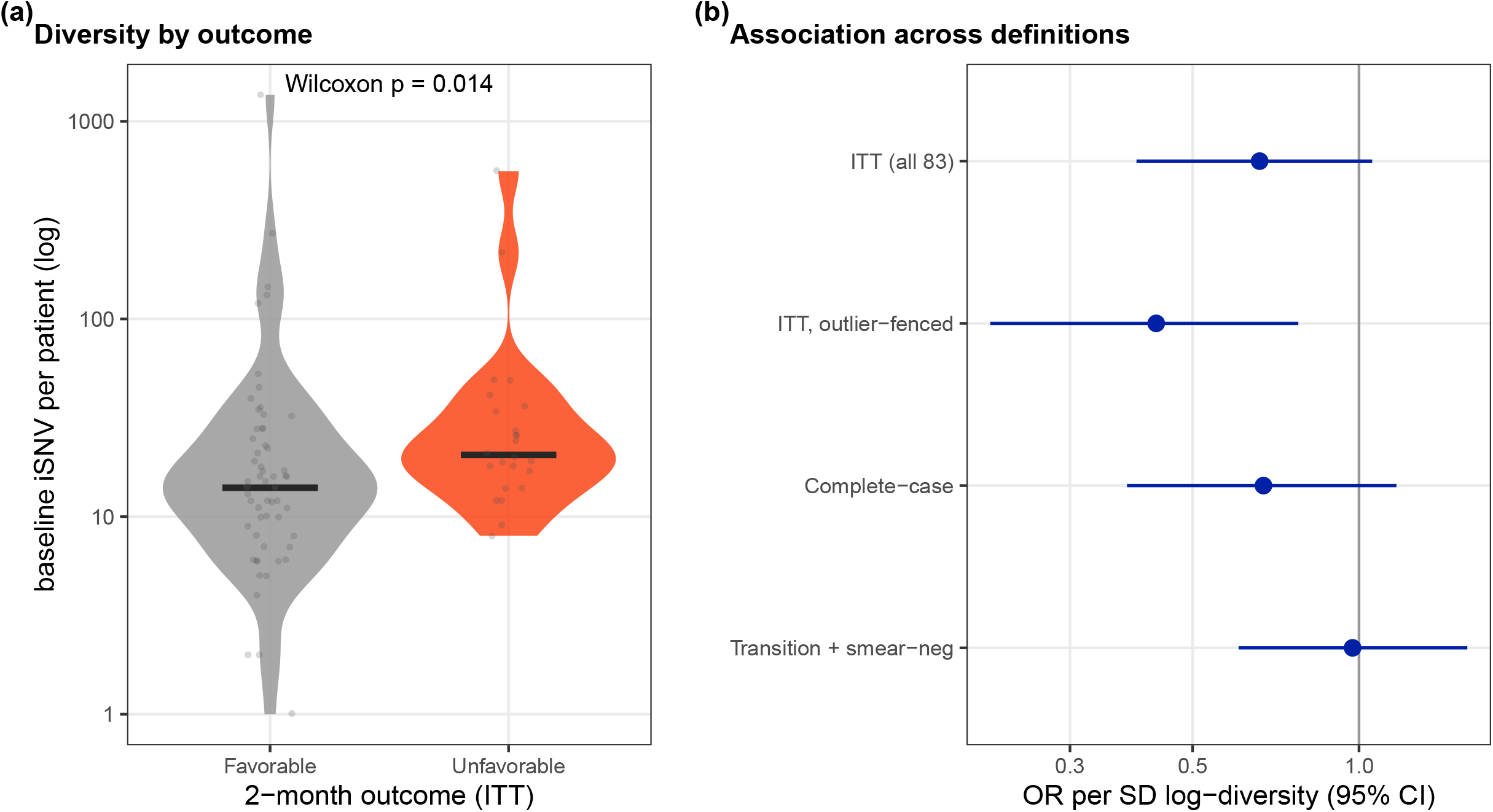
Baseline within-host diversity and 2-month treatment outcome. (a) Baseline iSNV burden by intention-to-treat (ITT) 2-month outcome (favorable = transition to continuation phase; unfavorable = failure, death, still-intensive, second-line, MDR, or loss to follow-up; 59 favorable, 24 unfavorable): higher in the unfavorable group (Wilcoxon p = 0.014). (b) Odds ratio per standard deviation (SD) of log baseline diversity for a favorable outcome across definitions (ITT, outlier-fenced ITT, complete-case, transition-plus-smear-negative); odds ratio (OR) below 1 means higher baseline diversity and worse outcome.

### Sampling and transmission scope

A single sputum sample undercaptured WHD. Spot and early morning specimens gave indistinguishable yields (unpaired p = 0.871; paired p = 0.147; **Figure S5a**), but paired specimens from the same patient each recovered only about two thirds of the combined variant union with limited overlap (fair one-versus-one comparison: spot 0.67, early morning 0.62, shared 0.23; **Figure S5b**); one specimen is a lower bound on within-host diversity. Shared variants were not transmission specific: they did not concentrate among patient pairs close by pathogen SNP distance or residence and linked and unlinked pairs did not differ in residential geographic distance (5.9 versus 5.0 km, p = 0.56; **Figure S6**). Given a random clinic sample of about 100 patients collected over twelve months, this null result reflects limited transmission signal in the sampling frame rather than evidence against transmission structure.

## Discussion

Within-host diversity in *M. tuberculosis* is increasingly used to infer transmission, detect emerging resistance, and predict treatment response, but WHD is measured inconsistently, and at the low frequencies where it lies, sequencing error and reference-mapping artifact are readily mistaken for biology. Setting the detection limit empirically from the per-site error process, mapping to the reconstructed complex ancestor[24], and masking above floor recurrent positions that conventional filters retain removed about a fifth of candidate calls and approximately halved the median per-patient burden. This deflation is consequential: a within-host estimate is only as reliable as the calls that underlie it, and halving the burden estimate is enough to alter qualitative conclusions. The modest corrected burden, a median of 16 minor variants per patient, is consistent with recent reports of within-host diversity below earlier estimates[15].

We report that WHD is elevated in *M. africanum* relative to L4. Because *africanum* is the lineage most divergent from an L4-derived reference[24], reference bias is the leading alternative explanation, but the contrast persists across masking thresholds and *africanum* calls were cleaner than L4 calls on the artifact-sensitive metrics, the opposite of what lineage-specific reference bias produces. The elevation is therefore predominantly biological, with a bounded residual at paralog and repeat loci that only an *africanum*-specific reference would fully resolve. The magnitude of the association, though not the direction, is uncertain at thirteen *africanum* patients. A lineage difference in WHD, in a West African cohort where *africanum* is common[7], cautions against generalizing L4-centric within-host expectations.

Within-host evolution over the first two months of treatment was dominated by drift and clearance rather than selection. Burden was stable while the underlying variants turned over, most sites were lost, and lost sites overwhelmingly cleared. The small fraction of sites that swept to fixation are better read as compartment or subclone replacement rather than as independent adaptive sweeps. This is consistent with prior descriptions of within-host dynamics under treatment[6,14] and argues against a strong directional signal in short-course, drug-susceptible disease.

Baseline within-host diversity was higher in patients with an unfavorable two-month outcome, the direction reported elsewhere[15]. Unlike the host associations, which disappeared under an outlier fence, this association strengthened when extreme-burden patients were removed, indicating that it is less susceptible to outliers. The association is nonetheless modest and underpowered: twenty-four unfavorable events, a full-sample logistic interval that includes the null, and no association when the outcome is restricted to smear conversion, so it tracks the clinical trajectory of progression, failure, death, and loss to follow-up rather than bacteriology alone. The finding motivates a larger, prospectively powered test. Baseline host and clinical features did not robustly explain burden after fencing outliers and shared within-host variants did not concentrate among transmission-plausible pairs. The transmission null carries little weight against transmission structure in principle: a random clinic sample of roughly one hundred patients over twelve months contains little transmission signal, and the result bounds what these data can support.

Our study has several limitations. Sequencing was deep short-read rather than long-read[37]; deep short-read data remain the public-health standard, and the detection limit and masking were built to be conservative against its failure modes, but residual artifact at the most repetitive loci cannot be excluded and the *africanum*-specific reference bias is bounded rather than removed. The cohort is single-site and modest, the *africanum* and follow-up subsets are small, and the outcome analysis is underpowered. Follow-up ended at two months, precluding inference about later relapse. The substrate depends on analytic choices, the detection limit, the recurrence threshold, and the cluster criterion, and a different pipeline would place the signal-to-noise boundary somewhat differently, although the reported results are stable across the sensitivity analyses.

The contribution is an empirically calibrated, artifact-aware within-host substrate, with reproducibility assessed against biological replicates, applied in an under-represented West African, *africanum*-inclusive cohort. Measured this way, within-host *M. tuberculosis* diversity in early treatment is modest, lineage-patterned, drift-dominated, and neither host- nor transmission-structured, with a modest indication that higher baseline diversity accompanies worse two-month outcome.

## Contributors

MNS conceived and designed the study, acquired funding, developed the analysis software, performed the formal analysis and visualization, administered the project, supervised the work, and wrote the original draft. JAM and MAO led the investigation, provided study resources, curated the data, and contributed to project administration. JC developed bioinformatic analysis software. MAB, CD, EK, MZ, SMA, and HMG contributed to the investigation and data curation. All authors contributed to reviewing and editing the manuscript. All authors had full access to all data in the study and accept responsibility for the decision to submit for publication. MNS and JAM directly accessed and verified the underlying data reported in the manuscript.

## Acknowledgement

We thank the study participants and the clinical and laboratory staff at Korle Bu Teaching Hospital, Greater Accra Region, Ghana.

## Data availability

Sequence reads have been deposited in SRA under BioProject accession PRJNA1466981. Analysis code (the calibration, masking, and analysis pipeline) is available at https://github.com/mapouseraphin/k01-tb-genomics-paper1. Derived per-patient within-host diversity data underlying the figures are available on request. An earlier version of this work is available as a preprint on bioRxiv: https://doi.org/10.64898/2026.05.27.727477.

## Ethical standards

The study was approved by the Institutional Review Board of the University of Florida (IRB202003042) and the Institutional Review Board of Korle Bu Teaching Hospital, Accra, Ghana (IRB/000135/2020). Written informed consent was obtained from all participants.

## Use of artificial intelligence tools

The authors used OpenAI’s GPT 5.6 for copy-editing and language refinement, and Consensus (Consensus NLP, consensus.app) to support the literature review underpinning the Research in context panel. All scientific content, study design, analyses, results, and conclusions were developed and verified by the authors, who assume full responsibility for the manuscript.

## Supplementary Tables and Figures

**Table S1a.** Primary within-host variant calling procedure: caller, quality filters, and thresholds used to define analytic within-host variants.

| Component | Specification |
| --- | --- |
| Candidate window | minor-allele frequency in [0.02, 0.50]; biallelic SNVs only |
| Recurrent artifact mask | mask a position where a sub-2% minor allele reaches the depth-conditional floor in $\geq 3$ patients (resistance genes protected); 39 positions masked |
| Depth-conditional detection limit (LoD) | retain $AD_1 \geq a(DP)$; $a(DP)$ is the smallest alt-read count whose exceedance among monomorphic sites is ≤ target FPR 0.010; $DP$ 78/127/178/257/356/1703 $\rightarrow a$ 2/3/3/4/5/34 |
| Minor-allele strand-balance floor | strand balance $\geq 0.100$ (fixed conservative strand-bias filter; the weaker strand carries $\geq 10\%$ of minor-allele reads) |
| PE/PPE masking | PE/PPE and PE-PGRS loci masked in the primary analysis |
| Site depth floor | candidate positions with $DP < 50$ dropped (below the LoD calibration range) |
| Within-sample proximity de-clustering | drop any iSNV within 20 bp of another iSNV in the same sample |
| Cross-lineage artifact filter | remove a non-PE/PPE site only if iSNV-band in $\geq 50\%$ of patients across $\geq 2$ major lineages and not resistance |
| Estimation of data-derived parameters | artifact mask and the depth-conditional LoD estimated from the cohort's own monomorphic (fixed-class) sites |

**Table S1b.**
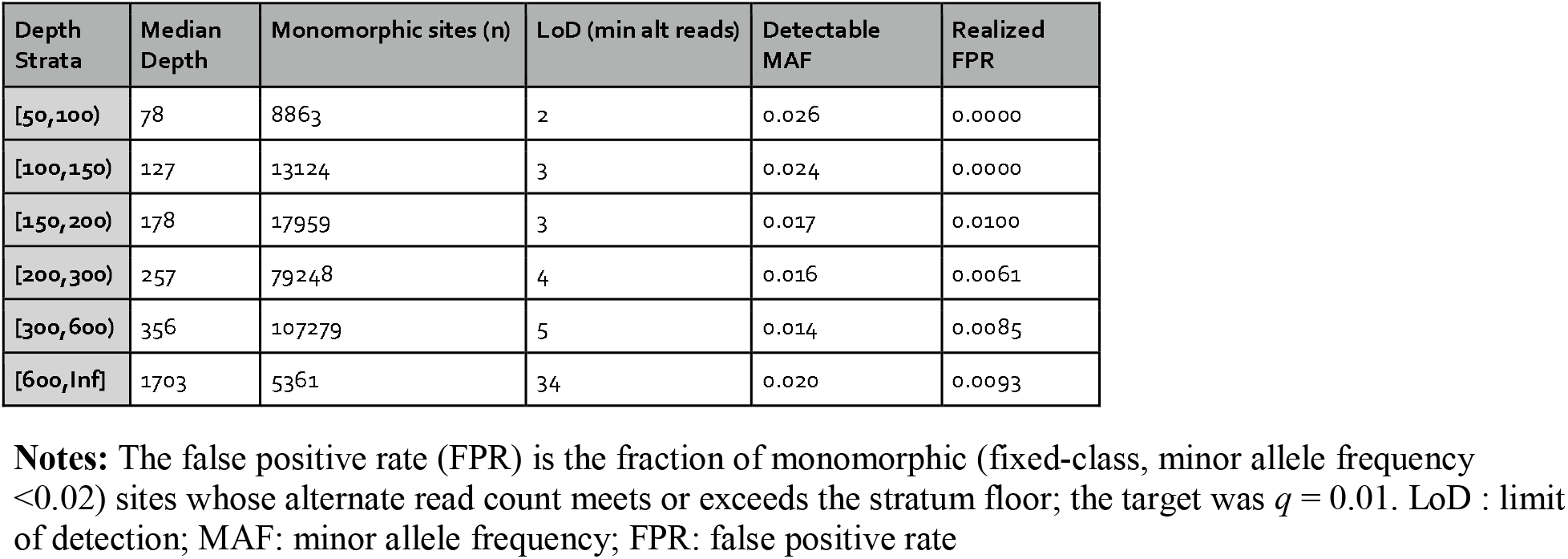
Depth-conditional limit of detection for within-host variant calling.

**Table S2.** Cross-lineage mixed-infection samples identified in the cohort.

| Sample | Strain | Patient |
| --- | --- | --- |
| KBTHo78-So.1 | lineage6.3.1;lineage4.6 | KBTHo78 |
| KBTHo78-So.2 | lineage6.3.1;lineage4.6 | KBTHo78 |
| KBTHo88-So.1 | lineage5.1.1;lineage1.1.3.2 | KBTHo88 |
| KBTHo88-So.2 | lineage5.1.1;lineage1.1.3.2 | KBTHo88 |
| KBTHo96-Eo | lineage6.3.1;lineage4.3.1 | KBTHo96 |
| KBTHo96-So.2 | lineage6.3.1;lineage4.3.1 | KBTHo96 |
| KBTH129-So.1 | lineage5.1.1;lineage4.6.2.2;lineage4.1.3 | KBTH129 |
| KBTHoo2-E1 | lineage4.9;lineage4.6.2.2 | KBTHoo2 |
| KBTHo23-So.1 | lineage4.9;lineage4.1 | KBTHo23 |
| KBTHo30-So.1 | lineage4.9;lineage4.6.5 | KBTHo30 |
| KBTHo30-So.2 | lineage4.9;lineage4.6.5 | KBTHo30 |
| KBTHo32-So.1 | lineage4.6.5;lineage4.6.2.2 | KBTHo32 |
| KBTHo32-So.2 | lineage4.9;lineage4.6.2 | KBTHo32 |
| KBTHo34-So.2 | lineage4.9;lineage4.6.5;lineage4.6.2.2 | KBTHo34 |
| KBTH129-Eo | lineage4.6.2.2;lineage4.1.3 | KBTH129 |
| KBTH129-So.2 | lineage5;lineage4.1 | KBTH129 |

**Table S3.** Candidate band recurrent positions masked as reference-mapping artifact (n = 45).

| Position | Gene | Patients | Lineages |
| --- | --- | --- | --- |
| 580772 | senX3-regX3 | 88 | 4 |
| 1644362 | Rv1458c-Rv1459c | 75 | 4 |
| 2266553 | Rv2020c | 73 | 4 |
| 3820545 | Rv3401-Rv3402c | 54 | 3 |
| 4120983 | Rv3680-whiB4 | 51 | 2 |
| 2631620 | plcA | 47 | 4 |
| 3131473 | Rv2823c | 47 | 4 |
| 2266442 | Rv2020c | 45 | 4 |
| 663419 | nrdZ-Rv0571c | 41 | 4 |
| 2945004 | PE_PGRS45-Rv2616 | 33 | 2 |
| 3478698 | moaA1 | 30 | 4 |
| 3478719 | moaA1 | 30 | 4 |
| 3478467 | moaA1 | 29 | 4 |
| 888774 | Rv0794c-Rv0795 | 25 | 4 |
| 2401825 | Rv2141c-leuU | 25 | 3 |
| 1612585 | Rv1435c | 23 | 2 |
| 1443428 | Rv1289-Rv1290c | 21 | 4 |
| 4359135 | espK | 21 | 4 |
| 3336587 | Rv2980-ddlA | 20 | 3 |
| 3478272 | moaA1 | 20 | 4 |
| 2266660 | Rv2020c | 19 | 3 |
| 3232759 | amt-ftsY | 18 | 2 |
| 3281238 | mas | 17 | 3 |
| 3566943 | Rv3196A-Rv3197 | 17 | 3 |
| 3820407 | Rv3401-Rv3402c | 17 | 4 |
| 1224340 | phoH2-Rv1096 | 16 | 3 |
| 3707602 | sdhB-vapC44 | 16 | 3 |
| 4341402 | espE | 16 | 3 |
| 1168720 | Rv1046c | 15 | 1 |
| 3372612 | ligA | 15 | 3 |
| 4083327 | Rv3645 | 15 | 2 |
| 4199132 | serX | 15 | 3 |
| 745989 | Rv0648 | 14 | 2 |
| 3336646 | Rv2980-ddlA | 14 | 2 |
| 802501 | rplW-rplB | 13 | 2 |
| 4100844 | MTS2823 | 12 | 2 |
| 38991 | fadD34-Rv0036c | 11 | 2 |
| 600281 | Rv0508 | 11 | 2 |
| 1096470 | PE_PGRS18-mprA | 11 | 2 |
| 2470033 | Rv2205c | 11 | 3 |
| 935449 | lpqR | 10 | 1 |
| 1823342 | cydD-cydB | 10 | 1 |
| 2945167 | PE_PGRS45-Rv2616 | 10 | 1 |
| 3120151 | Rv2813-Rv2814c | 10 | 1 |
| 3691071 | lpdA-Rv3304 | 10 | 2 |

**Figure S1.**
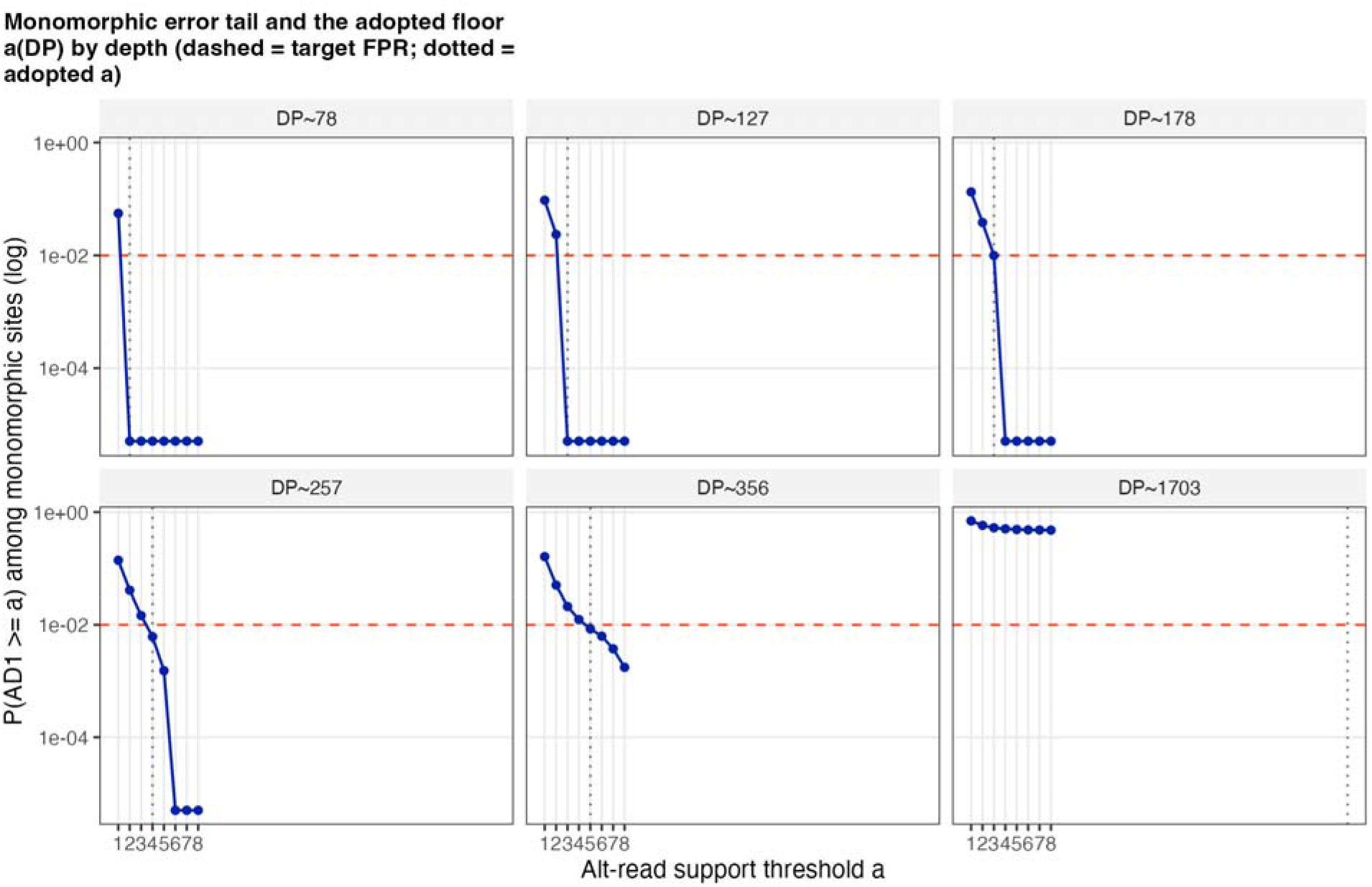
Monomorphic site error tail and the adopted detection floor, by depth. For each depth stratum, the probability that a monomorphic (fixed-class, minor allele frequency < 0.02) site reaches a given alternate read count, P(AD1 >= a), plotted against that count on a log scale. The dashed line marks the target false positive rate (q = 0.01); the dotted line marks the adopted floor a(DP), the smallest count at which exceedance falls to the target.

**Figure S2.**
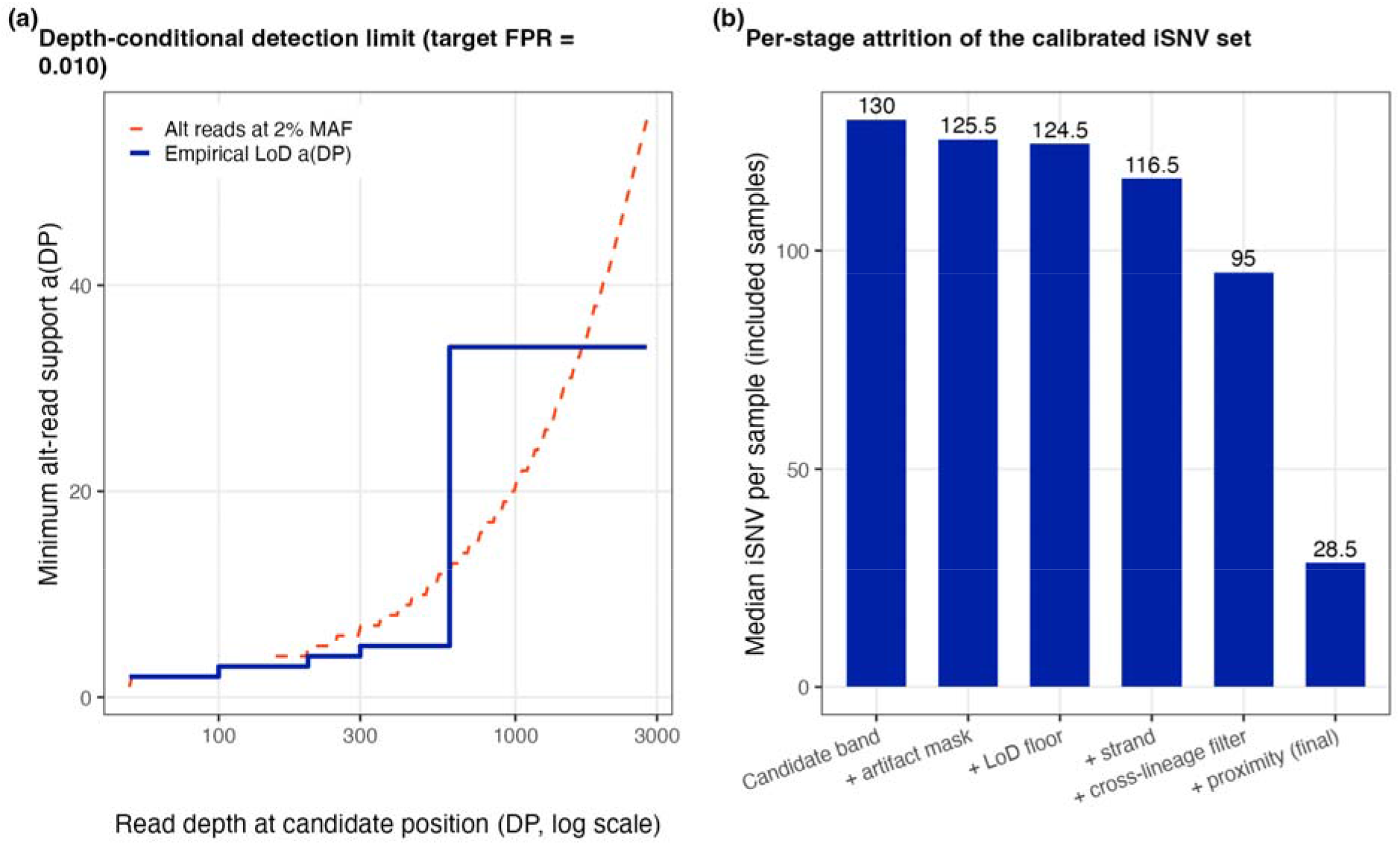
Detection floor and artifact-control attrition. (a) The depth-conditional empirical floor a(DP), drawn as a staircase over read depth (log scale; target false-positive rate q = 0.01). The dashed line marks the alternate-read count corresponding to a 2% minor allele: the floor tracks ∼2% at high depth and is count-limited (above 2%) at shallow depth. (b) Median within-host variants per included sample after each successive procedure stage, showing per-stage attrition of the calibrated set from the candidate band to the final analytic calls.

**Figure S3.**
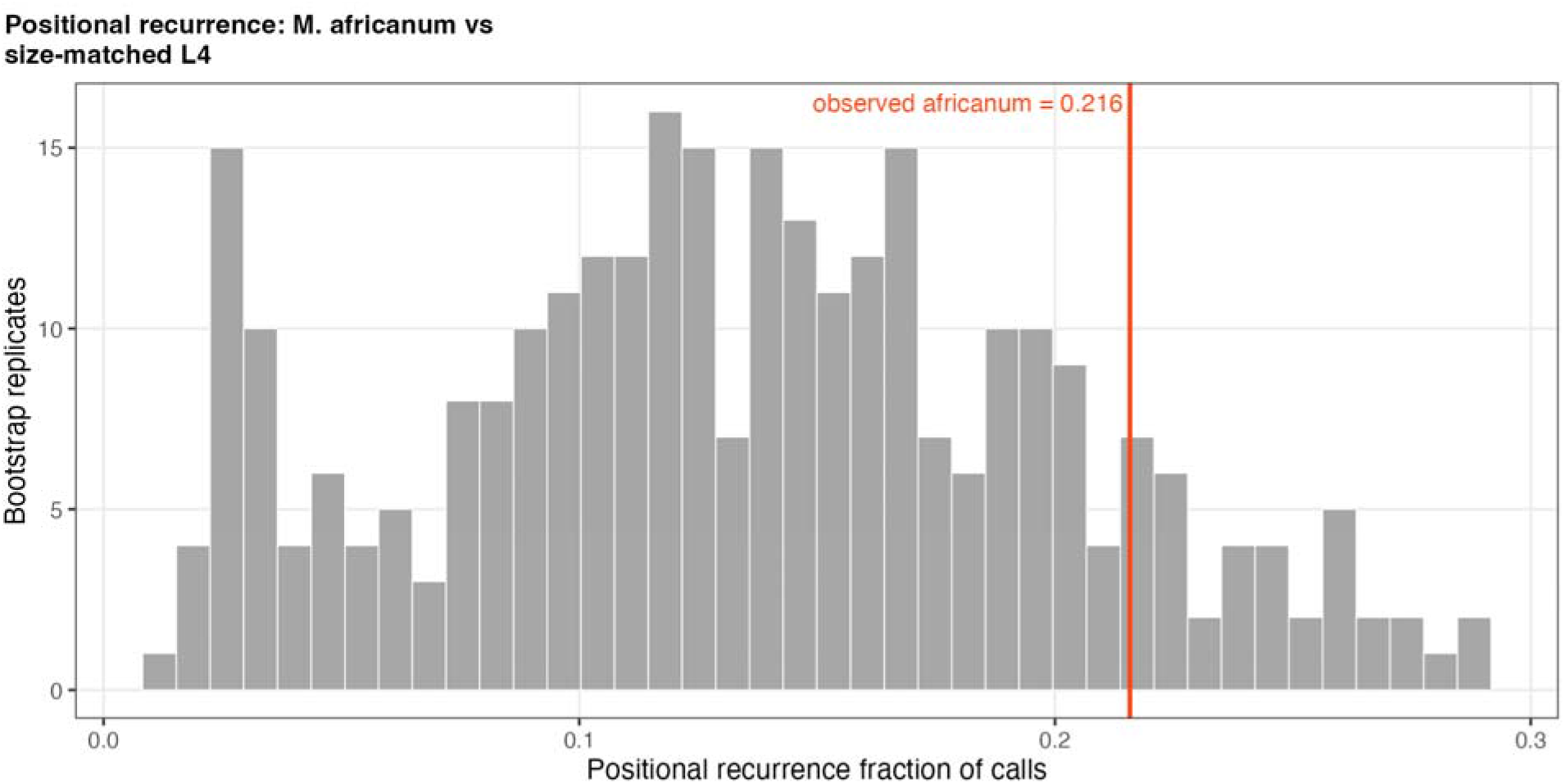
Positional recurrence: M. africanum versus size-matched lineage 4. Bootstrap distribution of the positional-recurrence fraction, the share of within-host variant calls falling at positions recurring across patients, for size-matched random draws of lineage 4 samples (grey). The orange line marks the observed value in *M. africanum. The africanum* elevation in within-host diversity is not explained by a higher burden of recurrent (artifact-prone) positions, which would have placed it in the upper tail of the L4 distribution.

**Figure S4.**
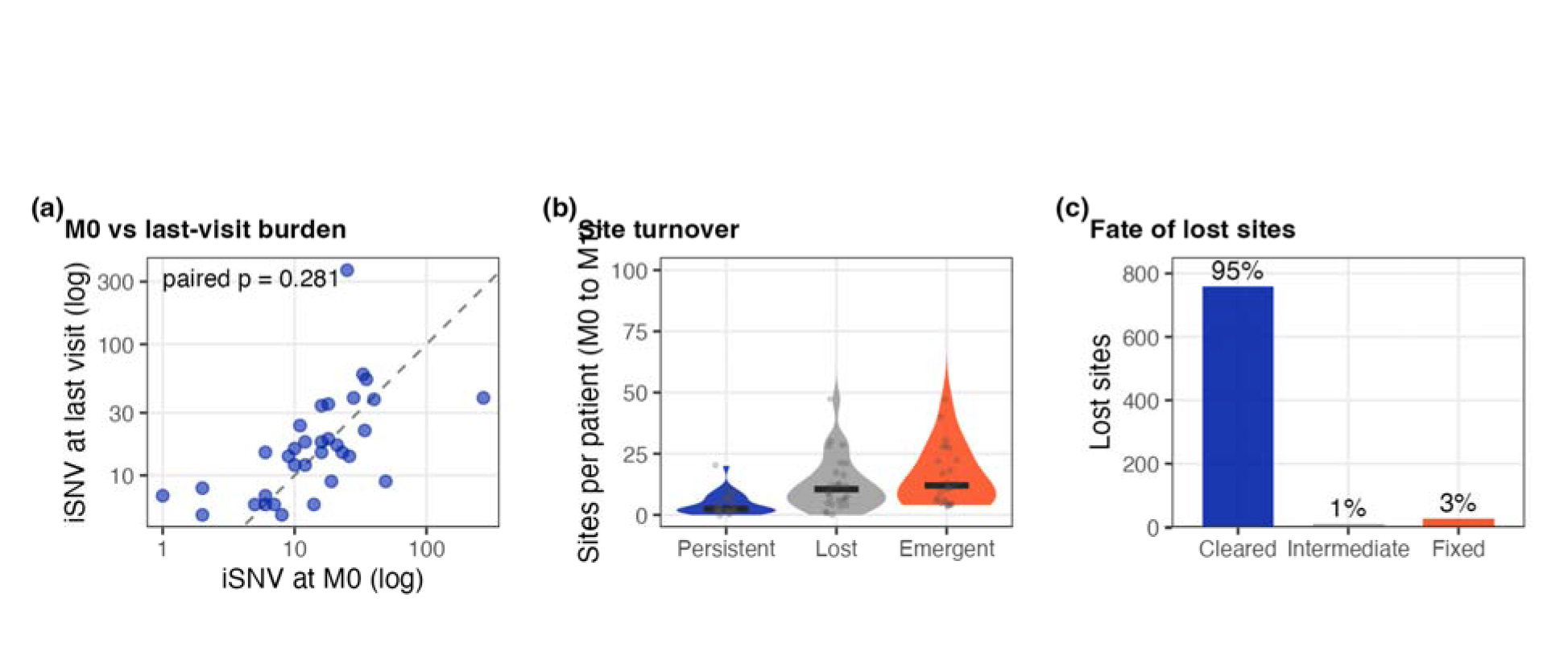
Longitudinal within-host dynamics over treatment. (a) Per-patient within-host burden at baseline (M0) versus the last available visit, on log axes; the dashed line is unity. Burden does not change systematically over treatment (paired p = 0.281). (b) Site turnover per patient from M0 to M2, partitioned into persistent, lost, and emergent variants; turnover is dominated by loss and emergence rather than persistence. (c) Fate of lost sites: 95% cleared to reference, 1% moved to intermediate frequency, and 3% fixed; loss reflects drift and clearance, not sweeps to fixation.

**Figure S5.**
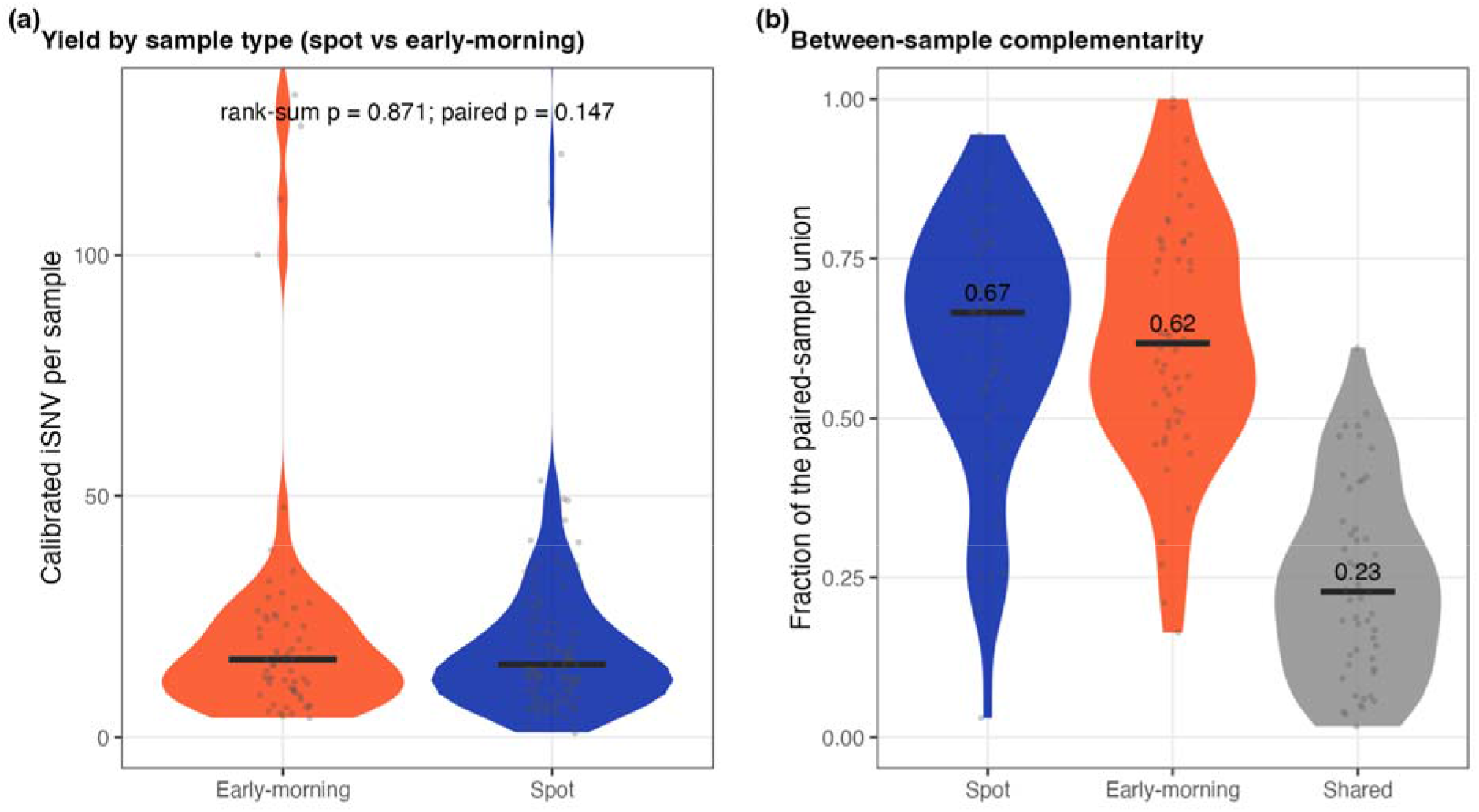
Single sputum sampling undercaptures within-host diversity. (a) Within-host variants per sample for early-morning versus spot sputum; yields are equivalent (rank-sum p = 0.871; paired p = 0.147). Neither specimen type is intrinsically richer. (b) Complementarity within paired specimens from the same patient: each specimen recovers only about two-thirds of the combined (paired-sample union) diversity (spot 0.67, early-morning 0.62), and just 0.23 of variants are shared by both. A single sputum therefore misses a substantial, non-overlapping fraction of within-host diversity. Black bars are medians.

**Figure S6.**
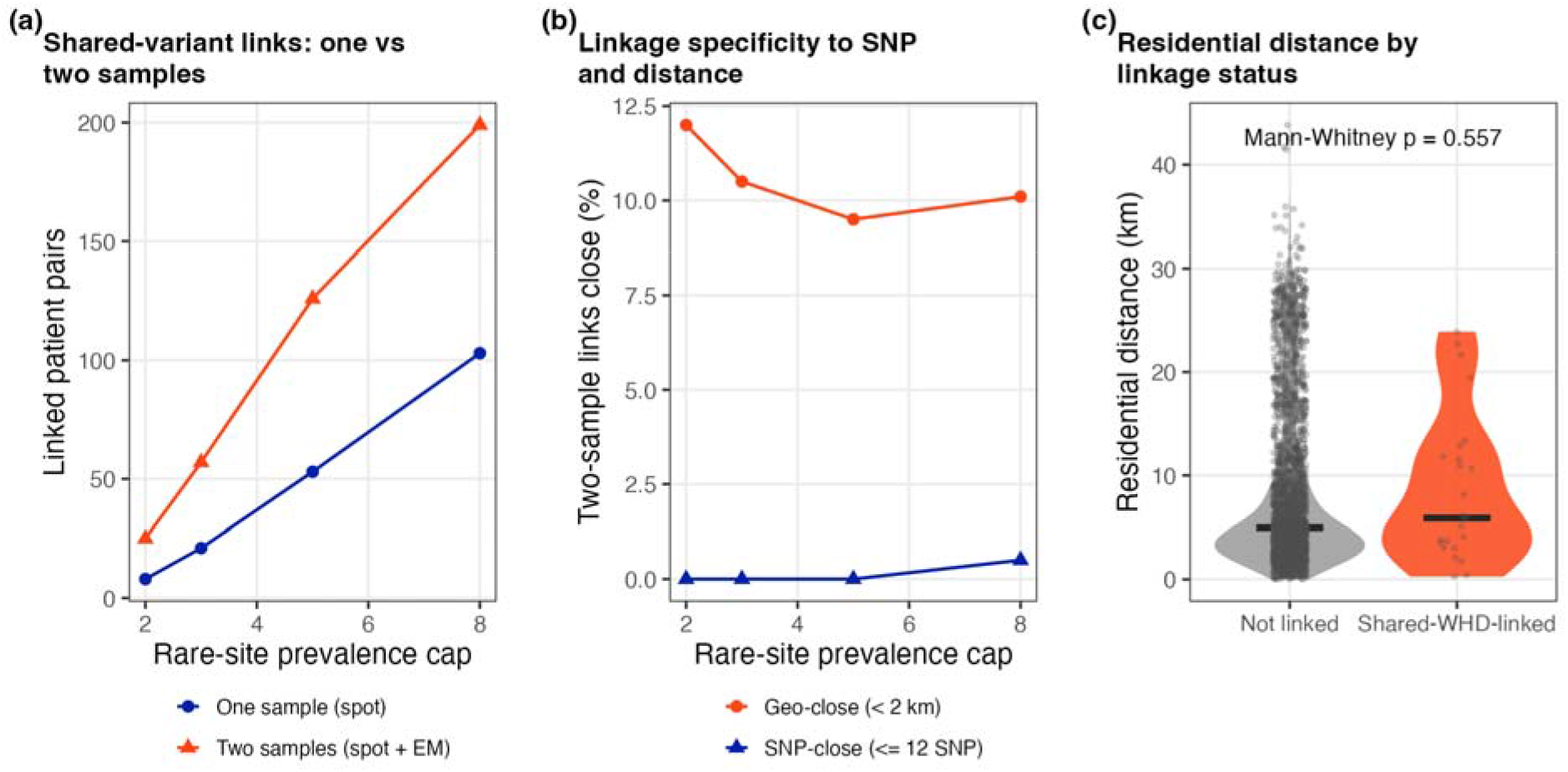
Shared within-host diversity do not mark transmission. (a) Number of patient pairs linked by shared rare within-host variants as a function of the rare-site prevalence cap, using one sample (spot) versus two samples (spot + early-morning) per patient. Adding a second specimen inflates the count of shared-variant links, indicating that sharing tracks sampling depth rather than a fixed biological signal. (b) Specificity of two-sample links: the percentage that are also genomically close (<= 12 consensus SNPs) or geographically close (< 2 km), across prevalence caps. Shared-variant links are almost never SNP-close, and only a small minority are geo-close. (c) Residential distance between patients who are shared-WHD-linked versus not; the distributions are indistinguishable (Mann-Whitney p = 0.557). Together, shared within-host variants show none of the genomic or spatial concordance expected of true transmission. Black bars are medians.

